# Systematic analysis of RNA-seq-based gene co-expression across multiple plants

**DOI:** 10.1101/139923

**Authors:** Hua Yu, Bingke Jiao, Chengzhi Liang

## Abstract

The complex cellular network was formed by the interacting gene modules. Building the high-quality RNA-seq-based Gene Co-expression Network (GCN) is critical for uncovering these modules and understanding the phenotypes of an organism. Here, we established and analyzed the RNA-seq-based GCNs in two monocot species rice and maize, and two eudicot species *Arabidopsis* and soybean, and subdivided them into co-expressed modules. Taking rice as an example, we associated these modules with biological functions and agronomic traits by enrichment analysis, and discovered a large number of conditin-specific or tissue-specific modules. In addition, we also explored the regulatory mechanism of the modules by enrichment of the known cis-elements, transcription factors and miRNA targets. Their coherent enrichment with the inferred functions of the modules revealed their synergistic effect on the gene expression regulation. Moreover, the comparative analysis of gene co-expression was performed to identify conserved and species-specific functional modules across 4 plant species. We discovered that the modules shared across 4 plants participate in the basic biological processes, whereas the species-specific modules were involved in the spatiotemporal-specific processes linking the genotypes to phenotypes. Our research provides the massive modules relating to the cellular activities and agronomic traits in several model and crop plant species.

## Introduction

The complex cellular network formed by the interacting macromolecules underlie an organism's phenotypes ^1-3^. Biomolecules are often thought to organize into interacting modules (functional building blocks) for completing a specific biological process ^4-6^. This standpoint is supported by the fact that many observable phenotypic variances are often not determined by a single gene but by a set of interacting genes ^7^. Systematic reconstructing a complete map of these interacting molecular modules are crucial for understanding an organism's genetic architecture underlying phenotypes.

Several methods have been developed to find functional gene modules by utilizing transcriptome data. Differential Expression (DE) analysis uses traditional statistical hypothesis testing-based approach, such as t-test, F-test, ANOVA or negative binomial test for assessing statistical significance of an observed expression change of each individual gene by comparing the between-conditions variation and within-condition variation, which can reveal the genes related to specific experimental conditions or sample types ^8-10^. However, differentially expressed genes are only a proxy for finding the key molecular modules related to our concerned biological questions because of highly dynamic transcriptome in different types of cells, tissues and experimental conditions ^11^. Complementary with the DE analysis, differential gene co-expression analysis aims to identify a group of differently co-expressed genes under two or more conditions, which has been applied to discern condition-specific gene co-regulation patterns ^12-15^. Differential co-expression analysis is especially effective in detecting biologically important genes that have less dramatic expression changes for certain conditions ^16,17^. Other than the two methods above, bi-clustering analysis is an approach that performs simultaneous clustering on genes and conditions across a wide range of transcriptome experiments. This method can discern the groups of genes that demonstrate similar expression patterns underlying the specific conditions but behave independently under other conditions. Though bi-clustering can identify a broad set of overlapping modules and thus present a global perspective on transcriptional network, genome-wide application of this approach is generally hampered by its inherent high computational complexity ^18^. Gene co-expression meta-analysis is another powerful method, which adopted the all experimental conditions to build co-expression network. When compared with bi-clustering analysis ^19-23^, its simplicity make it a powerful tool for identifying transcriptional modules.

In this study, using the ensemble pipeline used to build the rice RNA-seq-based Gene Co-expression Network (GCN) (unpublished method, under review), we further built the RNA-seq-based GCN in one monocot species of maize and two eudicot species of *Arabidopsis* and soybean and delineate them into co-expressed modules. Taking rice as an example, we associated the modules with biological functions and agronomic traits, and found a large number of condition-specific and tissue-specific modules. In addition, we also investigated the transcriptional regulatory mechanisms of modules by integrating known *cis*-element, transcription factors and miRNA targets. Moreover, we performed the comparative analysis of co-expressions across the 4 plant species to find the conserved and species-specific functional modules. Our research revealed the massive gene modules associating with the cellular activities and agronomic traits in several model and crop plant species, which provides a valuable data source for plant genetics research and breeding.

## Results

### Topological and biological properties of RNA-seq-based GCNs

The topological and biological properties of 4 RNA-seq-based GCNs built using the ensemble inference pipeline were analyzed. All these networks show the small-world characteristic with an average path length between any two nodes are smaller than 7 (Table S2). The distributions of node degrees obey the truncated power-laws where most nodes have a few co-expression partners with only a small ratio of hub nodes associating with a large number of partners (Fig.S1). We found that hub genes (with degree >200) were more functionally diversified than random ones in all four species (Wilcoxon rank sum test, *p*-value=8.46E-3 for *Arabidopsis*, *p*-value=6.23E-4 for rice, *p*-value=1.18E-7 for maize and p-value=2.20E-4 for soybean). This indicated that the hub genes of the co-expression networks might not be necessary to participate in central biological functions but provide the cross talks between different biological processes ^24^. On the other hand, we found that the likelihood of a gene to be essential increases with its degree, betweenness and closeness centricity, and they were more conserved across all plants (Fig.S2-S5). The negative correlation between the degrees (*K*) and the clustering coefficients (*C*) of genes revealed hierarchical and modular natures of these networks and the possible synergistic regulation of gene expression (Fig.S6) ^21^.

### Function and synergistic regulation of rice co-expression modules

One important feature of co-expression network is the modular structure, with genes sharing more connections within the module than between the modules. We adopted the Markov CLustering (MCL) method to obtain 772 gene co-expression modules (the number of genes > 5) in rice (Dataset 3). Of these modules, 771 modules are enriched in GO terms, pathways, protein functional domains or Tos17 mutant phenotypes. We found that the genes in co-expression modules shared more similar biological roles than the random selected genes (Wilcoxon rank sum test, *p*-value = 5.06E-07). Based on the enriched functions and gene expression patterns, we selected 12 gene modules participating in fundamental and condition-specific processes for further analysis (see Supplementary Text, Dataset 4, Dataset 5, Fig.S7-Fig.S10 for details). Among them, 5 gene modules are involved in photosynthesis; 4 modules are related to development of the reproductive organs, 2 modules were associated with cell cycle regulation and 2 modules were related to stress responses. For example, we found that two modules showing the pollen specific expression patterns (Fig.1) include a large amount of genes involving in the cell division, pollen germination, pollen tube growth and pollen sperm cell differentiation (Dataset 6).

**Fig. 1.**
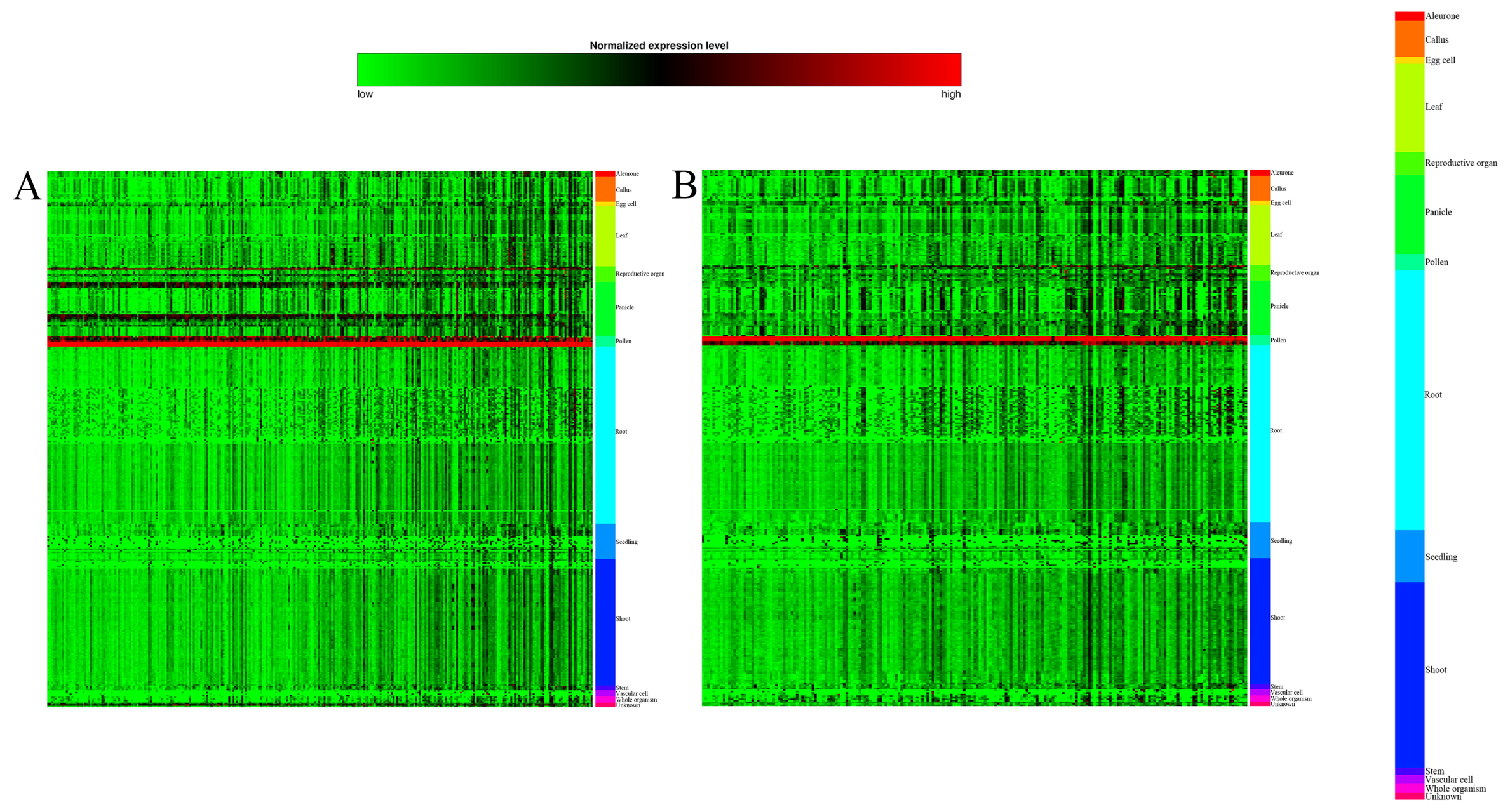
Co-expression modules related to the pollen development. A, Module #8, pollen-specific; B, Module #12, pollen-specific. The heatmap was produced by the VST dataset. In the heatmap, each row represents a sample, and each column represents a gene. The gene expression value was indicated by the color. The different colors of color bar on the right side represent the different types of tissues

The expression of a gene is often controlled by multiple factors such as transcription factors and miRNAs ^25^. Here, we further explored the regulation mechanisms of the co-expressed modules. We found known cis-elements in 770 modules, and found that 208 modules were enriched with targets of the same microRNAs and 291 modules were enriched with genes co-expressed with the common transcription factors. We also observed that the pairs of genes within co-expression modules, on average, have more common transcription factors and target genes of the same microRNAs than pairs of genes within random modules (Wilcoxon rank sum test, *p*-value=2.49E-28 for transcription factor and p-value=2.94E-28 for microRNA target). All these results suggested that the transcription factors or microRNAs tend to coordinately regulate targets sharing similar biological functions.

We found many examples in which the modules were simultaneously regulated by multi-factors. The most obvious example is the Module #5 involved in cell cycle and floral organ development, whose genes were linked together by two TCP transcription factors (*LOC_Os11g07460* and *LOC_Os02g42380*), one CPP transcription factors (*LOC_Os03g43730*) and three miRNAs (*osa-miR396, osa-miR156* and *osa-miR529*) (Fig.2 and Dataset 7). *LOC_Os11g07460* was co-expressed with 69 genes, in which 34 genes were associated with cell cycle, cell proliferation, cell differentiation, floral organ development and other development processes. Similarly, *LOC_Os03g43730* was connected with 51 genes, 25 of these genes were associated with cell cycle and plant development processes. And *LOC_Os02g42380* was associated with 41 genes, in which 19 genes involving in cell division and floral organ development. We found that 22 genes were associated with at least two of three key transcription factors mentioned above; of these genes, 16 genes play important roles in cell division, cell proliferation, cell differentiation and floral organ development. Moreover, we found that 3 genes were linked to three transcription factors above simultaneously, i.e. *LOC_Os11g07460, LOC_Os02g42380* and *LOC_Os03g43730*). These three genes were involved in flower morphogenesis, post-embryonic development and meristem growth. In addition to the transcription factors, the target genes of osa-miRNA156, osa-miRNA396 and osa-miRNA529 were also captured and enriched in the same module. Two of these three miRNAs (osa-miRNA156 and osa-miRNA396) play important roles in the cell division and organ development of the *Arabidopsis thaliana* ^26-28^. The common target *LOC_Os08g39890* of osa-miRNA156 and osa-miRNA529 was co-expressed with *LOC_Os07g03250*, which was related to the reproduction and development processes and was linked to two key transcription factors described above *(LOC_Os03g43730* and *LOC_Os11g07460)* and one MADS-box family gene. These results showed that synergistic regulation of co-expressed modules by multiple transcription factors and miRNAs.

**Fig. 2.**
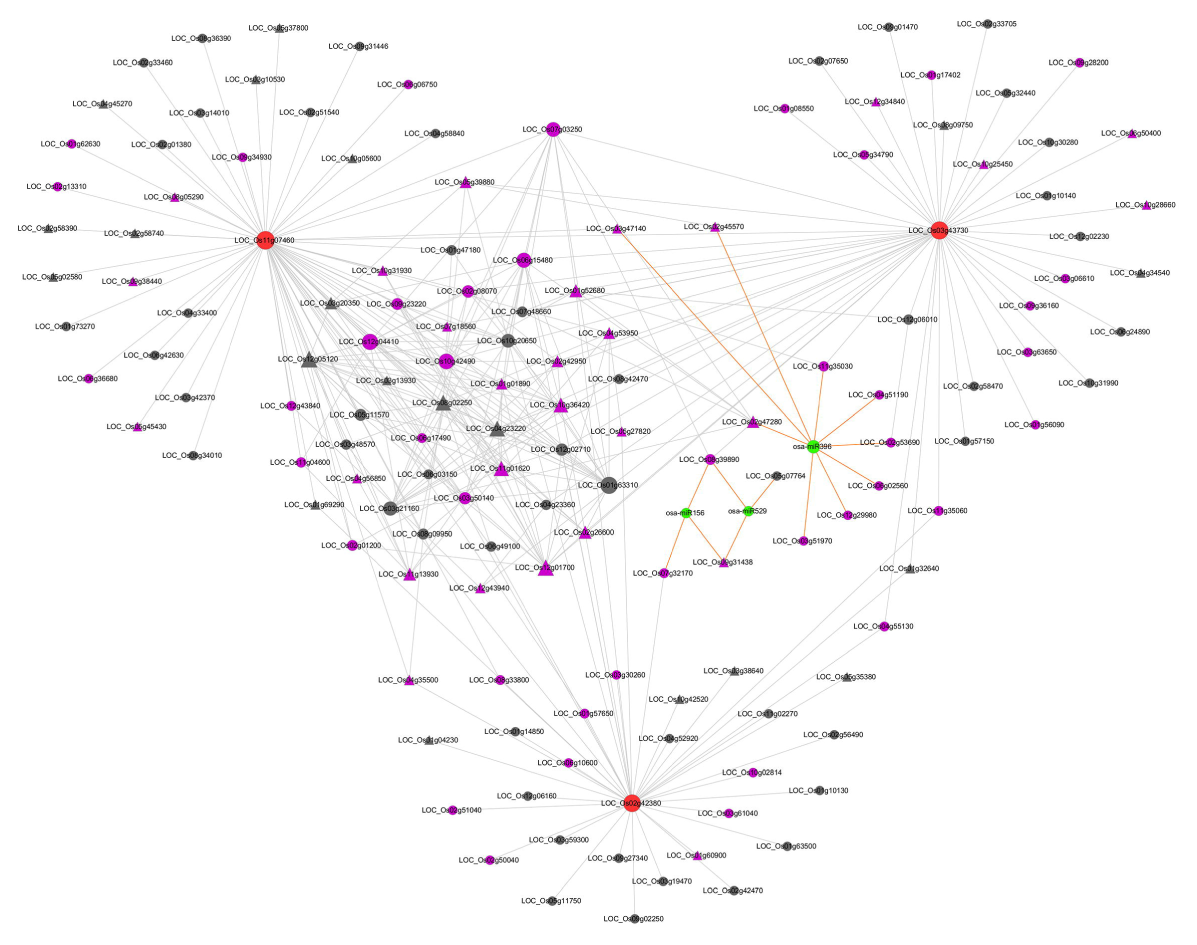
The synergistic regulation of Module #5 by multi-factors. Brown nodes indicate transcription factors; Green nodes denote miRNA; Pink nodes represent the genes involving in cell cycle, flower development or other development processes; Grey nodes indicate that the genes are function unknown or annotated with irrelevant functions; Triangle nodes denote the genes containing the known consensus motif of WTTSSCSS related to cell cycle. The size of node is proportional to the number of connected genes. For demonstration purpose, for except the co-expression links related to transcription factors (brown nodes)/miRNA (green nodes), we only showed the connections with confidence score larger than 0.2.

Furthermore, for the 12 modules mentioned above, we observed the strong coherence among the enriched transcription factors; known motifs and the enriched functions of modules (see Supplementary Text and Dataset 4 for details). For example, we found that two DBB transcription factors, 3 G2-like transcription factors and 5 CO-like transcription factors were tightly co-expressed with genes of photosynthesis modules (Table 1). In addition, 18 known cis-regulatory elements involving in light regulation are also enriched in these modules (Table 2). In another instance, we observed that the TCP, CPP and E2F/DP family of transcription factors are strongly linked to cell cycle modules (Table 2). And the known cell cycle motifs of E2FCONSENSUS, E2FAT and E2FANTRNR are also enriched (Table 2). For the pollen-specific modules, M-type transcription factors are tightly linked, and three known cell cycle *cis*-elements E2FCONSENSUS, E2FANTRNR and E2FAT were enriched (Table 1 and Table 2). In terms of stress response modules, WRKY, MYB, NAC and ERF transcription factors are linked with them. And three known stress response elements WBOXATNPR1, MYB1AT and ELRECOREPCRP1 are enriched (see Table 1 and Table 2). The less prevalent miRNA target enrichment in modules indicates that the biological functions of miRNAs and their target genes were diversified under evolution ^29^. Though the enrichment of miRNA targets in modules is infrequent, the roles of miRNAs and their targets can be inferred by the enriched functions of modules.

**Table 1.** The representative results of enriched transcription factors for the 12 rice co-expression modules

| Module ID/Function category | Gene ID | TF family | P-value |
| --- | --- | --- | --- |
| 1/stress response | LOC_Os08g09800 | WRKY | 6.16078E-94 |
| 1/stress response | LOC_Os08g09810 | WRKY | 1.22075E-89 |
| 1/stress response | LOC_Os02g15340 | NAC | 1.00472E-24 |
| 1/stress response | LOC_Os04g44670 | ERF | 1.51304E-23 |
| 1/stress response | LOC_Os04g43560 | NAC | 2.43893E-22 |
| 1/stress response | LOC_Os11g47460 | MYB | 2.64376E-66 |
| 1/stress response | LOC_Os09g26170 | MYB | 3.28823E-63 |
| 4/photosynthesis | LOC_Os04g42020 | CO-like | 7.87098E-35 |
| 4/photosynthesis | LOC_Os09g06464 | CO-like | 1.52885E-08 |
| 5/cell cycle | LOC_Os11g07460 | TCP | 4.0215E-72 |
| 5/cell cycle | LOC_Os03g43730 | CPP | 9.34839E-50 |
| 5/cell cycle | LOC_Os02g42380 | TCP | 1.40099E-44 |
| 5/cell cycle | LOC_Os02g42950 | YABBY | 1.32803E-93 |
| 5/cell cycle | LOC_Os01g52680 | MIKC | 2.45616E-84 |
| 5/cell cycle | LOC_Os07g32170 | SBP | 8.16666E-84 |
| 5/cell cycle | LOC_Os06g44860 | SBP | 2.70844E-57 |
| 5/cell cycle | LOC_Os02g08070 | SBP | 3.34117E-39 |
| 5/cell cycle | LOC_Os08g39890 | SBP | 3.94972E-38 |
| 5/cell cycle | LOC_Os09g31438 | SBP | 1.62399E-28 |
| 5/cell cycle | LOC_Os06g06750 | MIKC | 6.26973E-23 |
| 7/photosynthesis | LOC_Os04g41560 | DBB | 1.47199E-55 |
| 7/photosynthesis | LOC_Os06g24070 | G2-like | 0.002653096 |
| 7/photosynthesis | LOC_Os06g44450 | CO-like | 1.34414E-14 |
| 9/photosynthesis | LOC_Os07g48596 | G2-like | 0.001566656 |
| 10/cell cycle | LOC_Os06g13670 | E2F/DP | 2.16361E-13 |
| 10/cell cycle | LOC_Os02g50630 | E2F/DP | 2.25324E-11 |
| 10/cell cycle | LOC_Os12g41230 | CPP | 0.00126451 |
| 13/photosynthesis | LOC_Os02g39360 | DBB | 0.002057518 |
| 13/photosynthesis | LOC_Os12g01490 | G2-like | 4.90549E-15 |
| 13/photosynthesis | LOC_Os06g15330 | CO-like | 4.51511E-21 |
| 13/photosynthesis | LOC_Os02g39710 | CO-like | 2.35714E-08 |
| 15/stress response | LOC_Os07g22730 | ERF | 1.37837E-42 |
| 15/stress response | LOC_Os10g33810 | MYB | 2.46562E-13 |

**Table 2.**
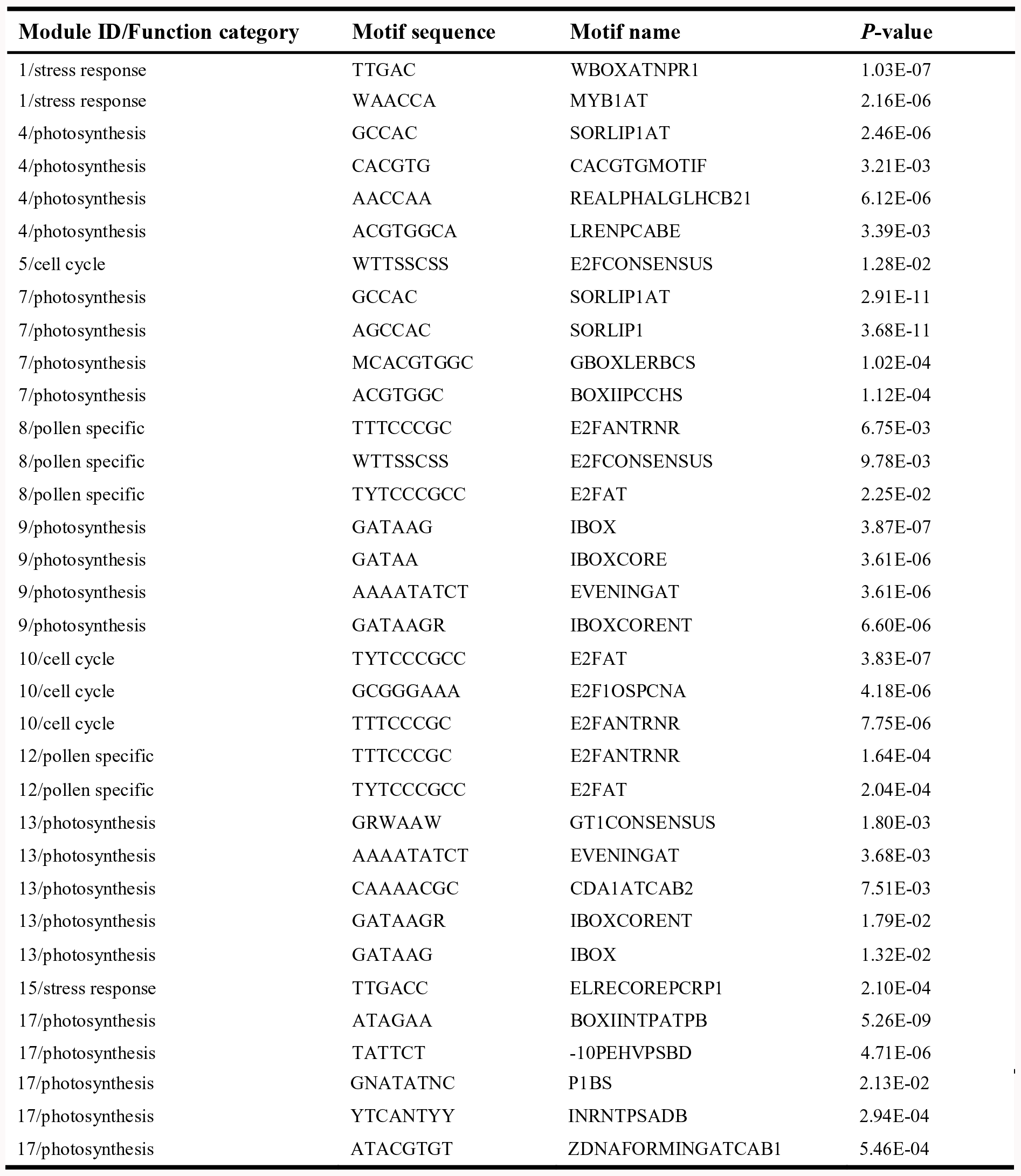
The representative results of enriched known cis-regulatory motifs for the 12 rice co-expression modules

### Co-expression modules controlling rice important agronomic traits

We asked whether the genes associating with common agronomic traits were placed and enriched in the co-expression modules. Interestingly, we found genes relating to the same agronomic traits were co-placed in the common modules (Table 3), which is consistent in functions with the agronomic trait. Firstly, it is expected that Module #7 whose genes were enriched in the agronomic traits of source activity. Secondly, Module #5 and Module #10 (modules participating in cell cycle) contain a large number genes relating to the agronomic traits of sterility and dwarf. Thirdly, genes associated with the agronomic trait of panicle flower were enriched in the Module #30. In addition, we also found that both Module #1 and Module #6 (containing the large number of pathogenesis-related transcription factors) whose genes were enriched in the agronomic traits relating to various resistances. Interestingly, we observed a module related to physiological trait of eating quality, and genes in this module were involved in starch biosynthesis, which is consistent with the fact that the component and molecular structure of starch are correlated with rice eating quality ^30,31^. These obtained results suggested that genes controlling the same agronomic traits were intrinsically clustered together in the network. According to the ranking of the number of links with known agronomic trait genes, we selected top 10 candidate biochemical function known and unknown genes associated with the dwarf, source activity, sterility and eating quality from Module #5, Module #7, Module #10 and Module #33, respectively. Indeed, some of these genes are likely associated with the agronomic traits, according to their molecular functions, which can provide guidance for future molecular breeding (Dataset 8). Particularly, we also observed that two QTL/GWAS candidate genes of *LOC_Os07g10495* and *LOC_Os10g42299*, related to leaf length, width, perimeter and area were placed in Module #7 ^32^ As annotated in MSU project, these two genes are highly expressed in leaf and seedling relative to other tissues, and is the molecular components of plastid. Moreover, we also found that a QTL/GWAS candidate gene *LOC_Os02g37850* associated with the spikelet fertility control were located in Module #10, this gene was involved in cell cycle and highly expressed reproductive organs of pistil and inflorescence ^33^.

**Table 3.** The statistic table of agronomic traits whose genes were enriched in modules

| Module ID | Agronomic trait | # of agronomic trait genes contained in module | # of all agronomic trait genes contained in module | Enrichment fold | p-value |
| --- | --- | --- | --- | --- | --- |
| 1 | Other soil stress tolerance <sup>a</sup> | 19 | 30 | 5.36 | 1.44E-06 |
| 5 | Dwarf <sup>a</sup> | 15 | 30 | 1.97 | 1.22E-02 |
| 6 | Drought tolerance <sup>a</sup> | 6 | 29 | 4.97 | 7.24E-08 |
| 6 | Salinity tolerance <sup>a</sup> | 13 | 29 | 4.71 | 1.52E-06 |
| 6 | Cold tolerance <sup>a</sup> | 12 | 29 | 6.39 | 4.10E-05 |
| 7 | Source activity <sup>a</sup> | 9 | 30 | 7.01 | 1.38E-12 |
| 10 | Sterility <sup>a</sup> | 8 | 16 | 3.41 | 5.92E-03 |
| 30 | Panicle flower <sup>a</sup> | 6 | 13 | 5.04 | 9.33E-06 |
| 33 | Eating quality <sup>a</sup> | 12 | 7 | 14.08 | 4.55E-07 |

<sup>a</sup> represents the agronomic traits extracted from Q-TARO database and literatures

### Comparative analysis of co-expression networks across multiple plants

We performed a comparative analysis of gene co-expression networks across multiple plants to identify conserved and species-specific co-expressed functional modules across closely related or distant plant species. We first examined to what extent the co-expressions are conserved among species. Indeed, a significant proportion of the pairs of genes whose co-expressions are conserved between the different plants (see Supplementary Text, Table S3, Table S4, Table S5, Dataset 9 and Dataset 10 for details). As demonstrated in Fig. 3, we can observe that the co-expressions are more conservative within monocotyledons or dicotyledons than between monocotyledons and dicotyledons. In addition, using the co-expression neighbors-based inference ^34^, we also found that the predicted functions of orthologous genes between species are more consistent than the random control genes (see Table S6 for details).

**Fig. 3.**
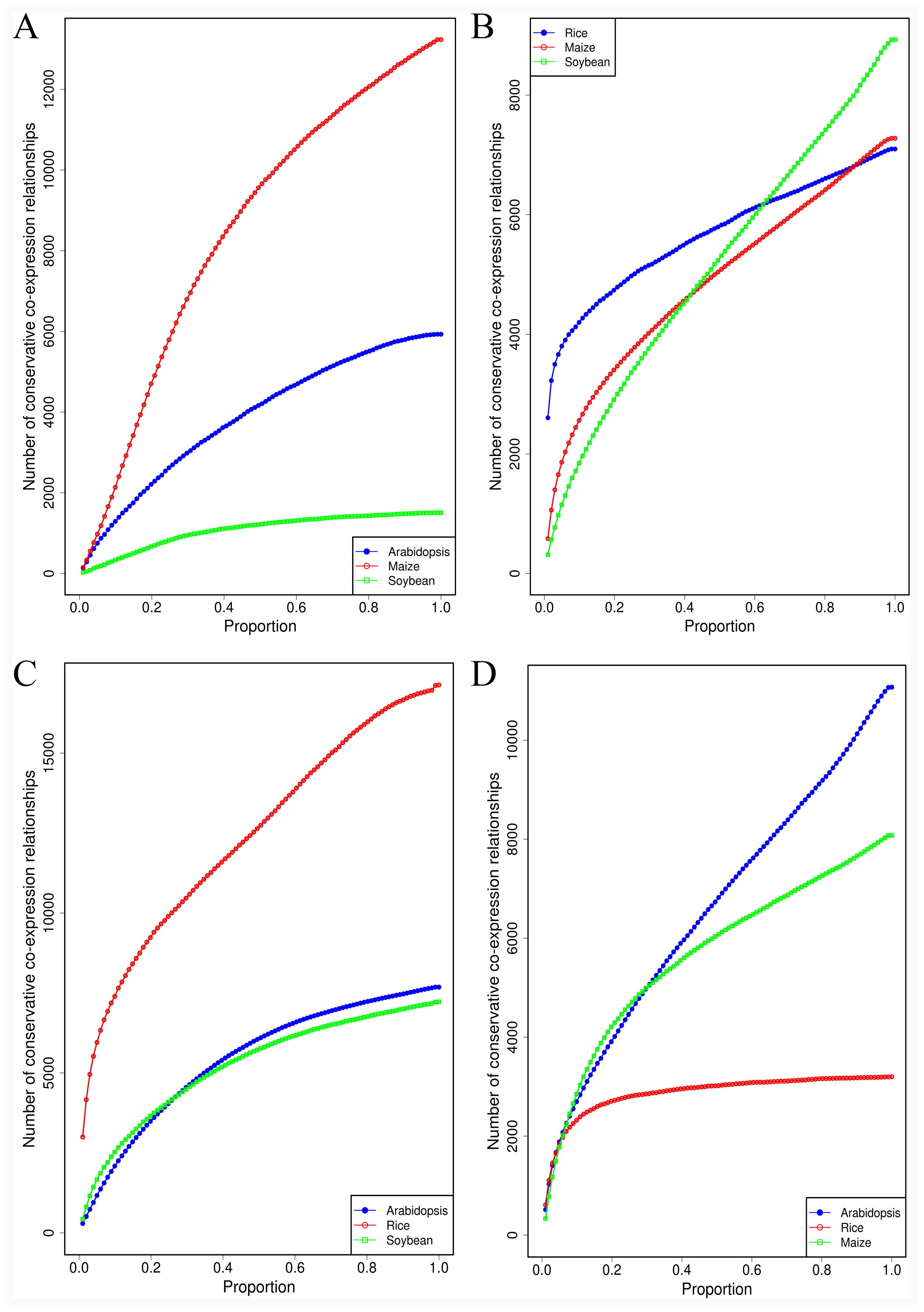
Number distributions of the conserved co-expression links between different plants at different proportion of co-expression links ranked by the confidence score

To analyze and compare the functional groups of these co-expression networks, we subdivided the network of each species into co-expressed functional modules based on co-expression link density and functional annotation similarity (for details, see Materials and Methods section). As a result, we here obtained 1396, 975, 1115 and 1065 modules for rice, *Arabidopsis*, maize and soybean, respectively. To assess the quality and reliability of obtained modules, we calculated for each real module the fraction of genes that own at least one homologue in a second species and compared with random modules. As expected, we found that the most modules have either significantly less or more homologous genes in other species than the random modules (Fig.4 and Table S7).

**Fig. 4.**
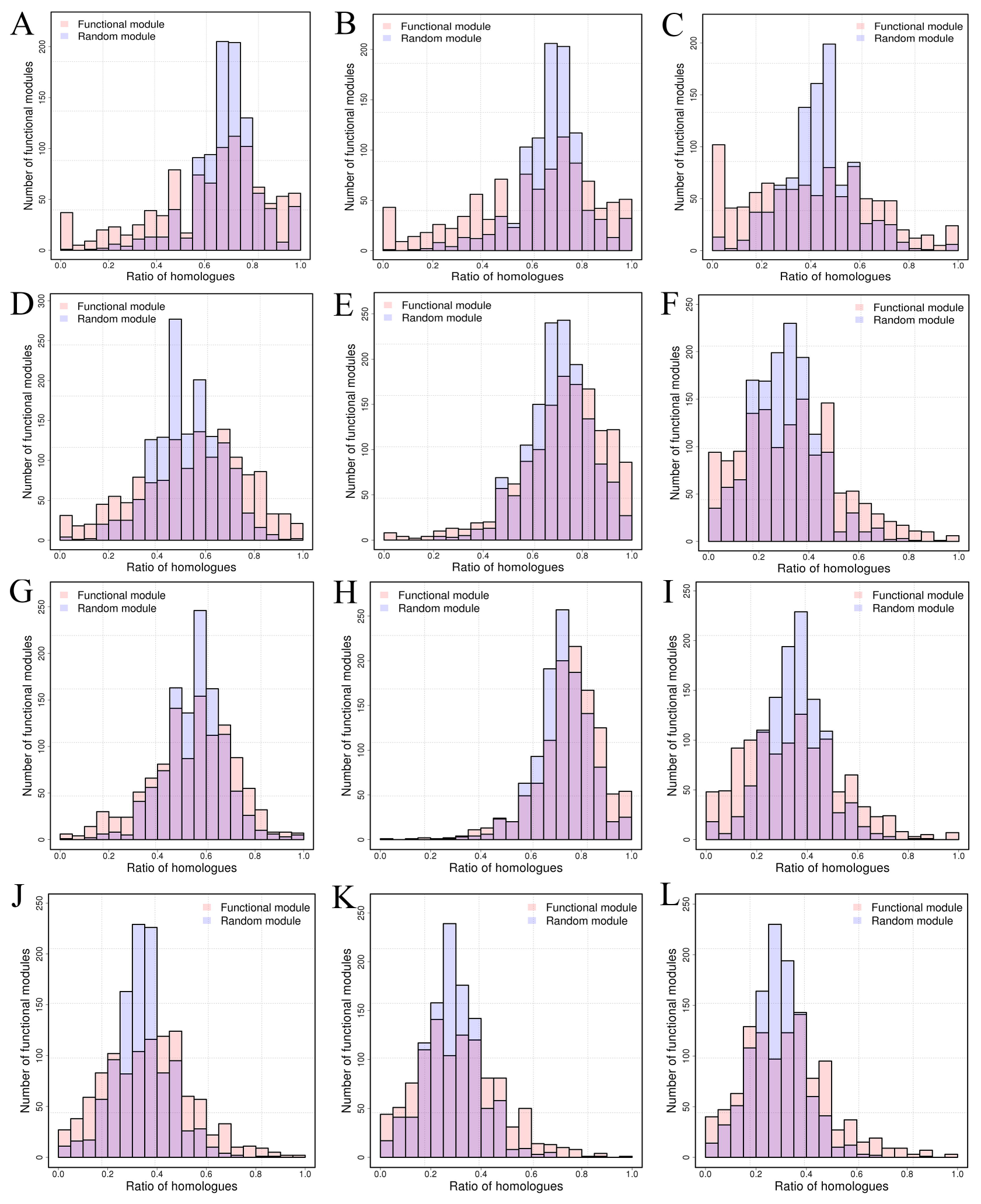
Histogram for the number of functional modules with a given fraction of genes possessing a homolog. Random module represents the random control distribution (preserving the same module size). A) *Arabidopsis* vs rice; B) *Arabidopsis* vs maize; C) *Arabidopsis* vs soybean; D) rice vs *Arabidopsis;* E) rice vs maize; F) rice vs soybean; G) maize vs *Arabidopsis;* H) maize vs rice; I) maize vs soybean; J) soybean vs *Arabidopsis;* K) soybean vs rice; L) soybean vs maize

We next focused on identifying the conserved functional modules sharing homologous genes across species and species-specific functional modules without homologous genes, based on the enrichment analysis of orthology relationships between genes for each combination of functional modules between four plants (see Materials and Methods for details). We defined a conserved module as one having homologous modules in at least one of the other species. Modules with no homologue module in all other plants are considered as species-specific. We identified 735 highly conserved modules among all four species (Dataset 11) and 2942 less conserved modules shared only by 3 or 2 plant species. Fig.5 demonstrated the common enriched GO terms of the functional modules within conserved modules. As expected, most of the common enriched GO terms are related to basic biological process, such as DNA replication, nucleosome assembly, RNA metabolic process, tricarboxylic acid cycle and cellulose synthetic process. We found that 62 best match of conserved module pairs (the percentage of homologous genes between two modules > 30%) with the common enriched GO terms, 48 module pairs enriched the same known motifs (length >= 6bp) in at least two species. In addition to the conservative modules, we also found 874 species-specific functional modules (Dataset 12). We observed that some species-specific functional modules whose genes are enriched in response to abiotic stresses, hormone stimuli and signal transduction, indicating a strong link between regulatory evolution and environmental adaptation. These results indicate that while a large amount of modules have been conserved under evolution, each species include more recently evolved modules linking genotype with phenotype.

**Fig. 5.**
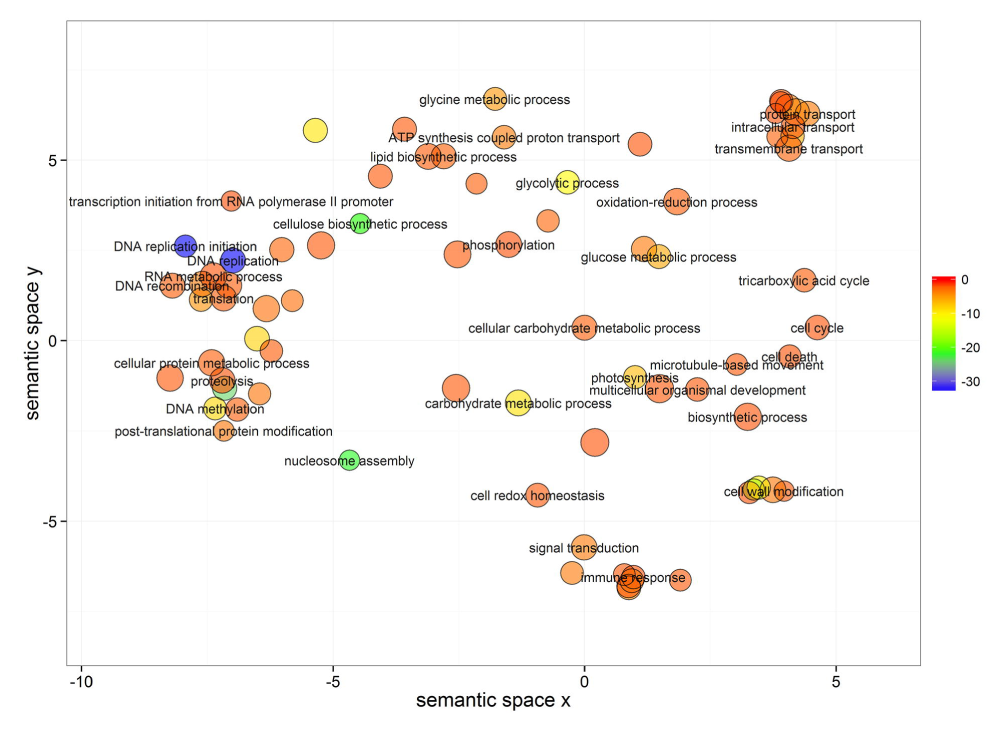
The common enriched GO terms of the functional modules within the conserved modules projected on the semantic space. The size of circle represents the gene number of GO term, and the color code indicates statistical significance

To describe the conserved and species-specific modules in details by examples, we further analyzed the inter-species conservation of 4 rice co-expression subnetworks as described in our previous literature (unpublished paper, under review) involving in cell wall metabolism, cell cycle, floral organ development and stress response process. We extended the single-species subnetworks into multi-species subnetworks by utilizing co-expression links within species and orthology relationships between genes across species. Note that we built the cross-species subnetwork involved in floral organ development process by expanding *AP3*-guide (an *Arabidopsis* homolog of rice *MADS16)* subnetwork, since genes in the MADS16-guide subnetwork have too few homologs in other plants. As expected, we observed that co-expressions are strongly conserved across all plants for cell wall metabolism and cell cycle processes (Fig.S11-S12). In contrast, the co-expressions in the subnetworks involving in stress response and flower development process were relatively less conserved between different plants (Fig.S13-S14).

## Discussion

In this study, we comparatively analyzed the high-quality RNA-seq-based gene co-expression networks and modules of 4 plant species: *Arabidopsis*, rice, maize and soybean, which were obtained by applying the pervious ensemble pipeline on the large amount of public available RNA-seq data (several hundred to more than one thousand samples for each plant species). The analysis of the topology properties of networks demonstrate that, for all these plants, the degree frequency distributions follow the truncated power-law; genes with high degree, betweenness and closeness tend to be essential and conserved between species; and network structure is highly modular. We also observed that the functionally related genes are often tightly connected together and the co-expression links are frequently conserved across the plant networks. The conserved and species-specific functional modules were identified using both the clustering analysis and orthology relationships enrichment of genes between different plant species. On one hand, the conserved modules across all plants provide an invaluable source for biological gene discovery and functional annotation transfer among different plants. On the other hand, a substantial ratio of modules has no significant conservation, indicating that novel genetics modules have been formed to accommodate the specific lifestyle and environment conditions. The similarity and difference of the modules between plants reveal the robustness and plasticity of gene regulatory networks. It is quite remarkable that some species-specific modules were enriched in the basic biological functions. For example, an *Arabidopsis* specific module of C8F1 plays important role in cell wall metabolism. It is interesting that the genes included in this module have no homologs in other plants. Similarly, five *Arabidopsis* chloroplast genome modules of C12F5, C12F7, C12F9, C12F11 and C12F12 involving in photosynthesis whose genes almost have no homologous genes in other plants. This result might be due to incorrect functional annotations of the genes and incomplete genome sequences.

Complementary with the co-expression neighborhood-based function prediction, the modules provide a valuable alternative for hypothesis-driven function inference of genes. The biological functions of uncharacterized genes in a given module could be inferred using the enriched functions of the modules. In addition, the conservative modules could be used to find functional analogous genome elements between species but their sequence have been diverged beyond recognition ^35^. Moreover, the conserved modules can also inherently remove the orthologs with similar sequence but not share similar biological functions, in contrast to sequence-based functional annotation ^21^.

Although co-expression between genes can be used to predict the gene functions, it is restricted to infer interactions where the regulators are co-expressed with their targets since it can only reveal the regulation of transcription level. Besides, co-expression network cannot also distinguish the regulators that are actually regulated a gene from ones that are simply co-expressed with a gene. We analyzed the regulatory mechanism of the modules by integrating the known motifs, transcription factor and microRNA targets. The outcomes demonstrated the strong agreements between the enriched known motifs, transcription factor, microRNAs and the enriched functions of modules. This agreement can be applied to infer the new regulatory interactions between the regulators and their targets.

## Materials and methods

### Experimental datasets

We downloaded the RNA-seq samples of rice, *Arabidopsis*, maize and soybean from the NCBI Sequence Read Archive (see Dataset 1 and 2 for details, accessed on May 29, 2014) using the same method as our previous study (reference). After the Sequence Read Archive (SRA) files were obtained, we transformed them into the FASTQ format using SRA Analysis Toolkit. The FASTQ sequencing reads files were firstly trimmed using Trimmomatic software (version 0.32) ^36^ with a parameter of the minimum read length at least 70% of the original size. Then, the fastq_quality_filter program included in FASTX Toolkit was used to further filtrate low quality reads, with the minimum quality score 10 and minimum percent of 50% bases that have a quality score larger than this cutoff value. The reads aligning and gene expression estimation were carried out by our previous analysis pipeline (reference). For *Arabidopsis*, maize and soybean, we used the TAIR10, Maizeb73v2 and Gmax_189 reference genomes for mapping and gene expression calculation. Gene Ontology (GO) annotations for all four plants were downloaded from the Plant GeneSet Enrichment Analysis Toolkit (PlantGSEA) ^40^. We extracted the biological pathways from three data sources including PlantGSEA, Gramene ^29^ and Plant Metabolic Network (PMN) database (http://pmn.plantcyc.org/). We obtained KEGG pathways from PlantGSEA for rice, *Arabidopsis* and soybean. Subsequently, we extracted the signaling and metabolic pathways in OryzaCyc, AraCyc, and SoyCyc databases from the PMN project data portal. With regard to maize, we integrated the pathway information retrieved from CornCyc database (contained in PMN project data portal) and MaizeCyc database (included in Gramene database). Besides, we also extracted the rice InterPro annotations from MSU Rice Genome Annotation Project website (http://rice.plantbiology.msu.edu/). The known agronomic trait genes were collected from the Q-TARO database ^41^ and literatures. Essential genes of *Arabidopsis* were retrieved from SeedGenes database ^42^. The known *cis*-regulatory motifs were extracted from both AGRIS and PLACE databases ^43,44^. Transcription factor families for all these plants were downloaded from the Plant Transcription Factor Database (PlantTFDB) ^45^. MicroRNAs and their related targets were collected from the Plant MicroRNA Target Expression database (PMTED) and Plant MicroRNA database (PMRD) ^46^. The orthologs between species were obtained by integrating the results of BLASTP alignment (with E-value < 1E-160), the predictions of OrthoMCL ^47^ and the known gene families provided in MSU Rice Genome Annotation Project.

### Module identification and enrichment analysis

A two-step decomposition procedure was adopted to identify the modular structure. We first divided the whole network into co-expression modules using an efficient graph clustering algorithm of Markov Clustering (MCL) with the default parameters (co-expression modules with the number of genes >= 5 were remained for subsequent analysis). Because the obtained co-expression modules might consist of hundreds of genes with numerous functional terms and multiple functional units, we carried out a second step to further subdivide the initial co-expression modules into non-redundant functional modules using functional annotation similarity clustering. Our clustering procedure adopted the Kappa statistics which is similar to the method used in ^48^, but with two important modifications. In details, a pair-wise Kappa *K* score was first calculated for each gene using the following equations:

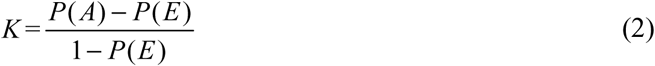

Where *P(A)* is the percentage agreement of functional terms between the gene pair, and *P(E)* represents the chance agreement. For rice, the GO, pathway, InterPro and Tos17 mutant phenotypes were combined as the functional terms. For *Arabidopsis*, maize and soybean, the GO and pathways were integrated as the functional terms. Based on the Kappa statistics, a seed cluster was formed for each gene by grouping it with all other genes with which it shares a *K* score greater than a given threshold. To obtain an appropriate threshold, we simulated 10000 background distributions of *K* score by randomly sampling 1000 genes from the genome space and used the average 95th percentile of these distributions as the *K* score threshold. Seed clusters with less than 3 genes were not considered. Also, seed clusters were only considered if 50% or more of the *K* scores between all group members were greater than the given threshold. Subsequently, the seed clusters were merged through an iterative process that exhaustively compared each cluster with every other group and merge any two that have more than 50% similarity. It continued until merging was no longer possible and the remaining clusters were treated as the functional modules. As many genes in the networks do not have the functional annotation, we adopted a procedure to assign these genes to the obtained functional modules. For each unannotated gene within a given co-expression module, we counted its connections with the genes of the functional modules derived from the co-expression module. Then, we selected the functional modules with the maximal links and moved the unannotated gene to these functional modules. This process continued until all unannotated genes were pushed to the functional modules. Note that we did not divide the co-expression modules with the number of annotated genes less than 3, and they were directly regarded as the functional modules. Functional modules were named after as follows: CxFy, where x is the number of co-expression module and y is the number of cluster. Note, for the very large co-expression modules cannot be subdivided into functional within 30 days using the in-house script, we further decomposed the sub-network composed of genes contained in each of these modules into smaller co-expression modules using different inflate parameters so that the co-expression modules can be effectively divided into functional modules.

The function, phenotype, known cis-regulatory motif and miRNA target enrichment of a module was calculated as the ratio of the relative occurrence in gene set of the module to the relative occurrence in the genome. To find known cis-regulatory motifs within each module, the promoter region (1kbp upstream from the transcription start site) of each gene in each module and entire genome was scanned for each known motifs. For each transcription factor, the enrichment of module was based on the ratio of the relative occurrences of genes co-expressed with the transcription factor between module and co-expression network. The statistical significance level was calculated using Fisher's exact test. The *p*-value smaller than 0.05 was regarded as enriched.

### Modules conservation analysis

To identify the conserved and specie-specific functional modules, the number of homologs pairs for the given two species was counted for each combination of the functional modules. The number of homologues pairs was then compared to the expected number based on the hypergeometric test,

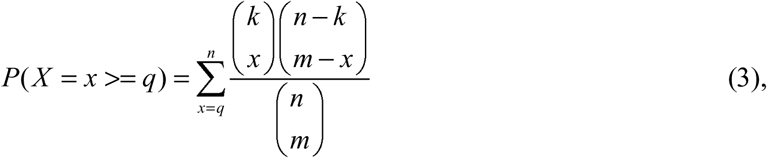

where *q* represented the number of orthologous pairs in combination of functional modules between the given two species, *k* was the total number of orthologous pairs between the given two species, *m* denoted the number of all possible gene pairs in the combination of functional modules between the given two species, and *n* presented the number of all possible gene pairs between the given two species. To as soon as possible obtain the true conserved modules and remove the false positives (e.g. produced by large plant gene families having many-to-many orthologs), the obtained *p*-values were further adjusted by the Benjamini-Hochberg correction for multiple hypotheses testing. Only the combinations with the q-value smaller than 0.05 were considered as homologous. Based on this, the conserved modules were defined as one having homologous modules in at least one of the other species. Modules with no homologue modules in all other plants are treated as species-specific. The enriched GO terms of modules were visualized using the tool REVIGO ^49^.

### Availability

The reconstructed RNA-seq-based co-expression networks and functional modules of 4 plant species can be freely downloaded at ftp://111111@ftp.mbkbase.org (username: 111111; password: 111111).

## Acknowledgements

This work is financially supported by The Strategic Priority Research Program of the Chinese Academy of Sciences (Grant No. XDA08020302). The funders had no role in study design, data collection and analysis, decision to publish, or preparation of the manuscript.

## Author Contributions

H.Y. conceived the original screening and research plans; H.Y. and C.Z.L. supervised the experiments; H.Y. performed the experiments and analyzed the data; B.K.J revised the paper; H.Y. conceived the project and wrote the article with contributions of all the authors; H.Y and C.Z.L supervised and complemented the writing.

